# Optical regulation of endogenous RhoA reveals selection of cellular responses by signal amplitude

**DOI:** 10.1101/2021.02.05.430013

**Authors:** Jeongmin Ju, Hae Nim Lee, Lin Ning, Hyunjoo Ryu, Xin X. Zhou, Hyeyeon Chun, Yong Woo Lee, Austin I. Lee-Richerson, Cherlhyun Jeong, Michael Z. Lin, Jihye Seong

## Abstract

How protein signaling networks respond to different input strengths is an important but poorly understood problem in cell biology. For example, the small GTPase RhoA regulates both focal adhesion (FA) growth or disassembly, but whether RhoA serves as a switch selecting between cellular outcomes, or if outcomes are simply modulated by additional factors in the cell, is not clear. Here, we develop a photoswitchable RhoA guanine exchange factor, psRhoGEF, to precisely control endogenous RhoA activity. We also develop a FRET-based biosensor to allow visualization of RhoA activity together with psRhoGEF control. Using these new optical tools, we discover that low levels of RhoA activation preferentially induce FA disassembly in a Src-dependent manner, while high levels induce both FA growth and disassembly in a ROCK-dependent manner. Thus, rheostatic control of RhoA activation with photoswitchable RhoGEF reveals that cells can use signal amplitude to produce multiple responses to a single biochemical signal.

## INTRODUCTION

A fundamental question in cellular computation is whether different activation levels of a single biochemical signal can be used to generate distinct functional outputs. For example, overexpression of individual members of the Rho family of small GTPases induces specific actin-based structures, with Cdc42 promoting filopodia formation and extension, Rac1 lamellipodia formation and extension, and RhoA actinomyosin contractility ^1^. However, experiments with fluorescent reporters have shown that, even within the same cell, individual GTPases are activated at sites and times of seemingly opposite morphological responses. For instance, in migrating cells, RhoA is activated both at the leading edge, which is extending, and at the lagging edge, which is retracting ^2–5^. Similarly, RhoA is required for focal adhesion (FA) growth in response to various extracellular stimuli ^1,6^, but has also been implicated in the reverse process of FA disassembly ^7,8^. Given its apparent involvement in the induction of opposing morphological changes, RhoA signaling to FAs could serve as a model system for understanding how a single regulatory protein can produce divergent downstream effects.

While mechanisms from RhoA to FA assembly have been extensively studied ^1,6^, much less is known about the mechanisms linking RhoA to FA disassembly. FA formation is a reliable static outcome of many extracellular stimuli, but FA disassembly has primarily been studied in the context of cell migration, when FAs are continuously assembled and disassembled ^6^. Indeed, localized FA disassembly allows cell regions to detach from their environment and thus is of essential importance in cell migration and cancer metastasis ^9^. RhoA has been hypothesized to drive lagging edge retraction based on biosensor imaging in migrating cells ^2,5^, but it is unclear whether RhoA is responsible for FA disassembly during this process, or how such a function would be reconciled with the better understood function of RhoA in FA growth.

Global loss-of-function techniques have been used to study RhoA effectors in FA disassembly, but it is difficult to dissociate primary from secondary phenotypes in such experiments. For example, dominant-negative constructs, RNAi, or chemical inhibitors of the mDia and ROCK families of RhoA effectors (hereafter referred to collectively as mDia and ROCK) produce trailing edge phenotypes in migrating cells, suggesting functions in FA disassembly ^7,8,10^. However, these global manipulations disrupt protein function throughout the cell over long periods of time, and mDia and ROCK also have well characterized roles in FA assembly and growth at the leading edge ^11–16^. Any observed deficits in FA disassembly in migrating cells upon interference with mDia or ROCK function could thus be secondary to earlier deficits in FA growth at the leading edge.

To clearly elucidate molecular mechanisms governing cell responses to specific protein activities, such as FA disassembly in response to RhoA, a method to activate a specific protein separately from other signaling pathways at a particular time would be useful. Here, we report the engineering of a photoswitchable activator of endogenous RhoA, psRhoGEF. We also develop a RhoA biosensor based on fluorescent resonance energy transfer (FRET) with LSSmOrange and mKate as a FRET pair to measure the RhoA activity induced by psRhoGEF without spectral overlapping. By applying different doses of light to psRhoGEF for rheostatic RhoA activation, we find that lower RhoA activity preferentially promotes FA disassembly while higher RhoA activity promotes both FA growth and disassembly. Interestingly, the switch to FA growth appears to involve suppression of Src, as Src activation at FAs occurs preferentially at lower levels of RhoA activation, and inhibition of Src-family kinases further promotes FA growth in response to RhoA activation. These results thus elucidate a specific role of Src in RhoA-induced FA disassembly, and explain how a biochemical signal can produce opposite outcomes depending on signal amplitude and context. Our results also demonstrate how new optical control of protein activity overcomes limitations of traditional methods in investigating complex cellular behaviors.

## RESULT

### Strategies for optical control of endogenous RhoGTPases

Previously two general strategies have been devised for optical control of Rho-family GTPase activity. The first strategy involves expressing optically controllable forms of the GTPase itself. For example, RhoA fused to cryptochrome 2 (CRY2) is clustered by blue light, resulting in its activation by unknown mechanisms ^17^. Alternatively, RhoA can be sequestered to mitochondria via a protein-protein interaction that occurs in the dark but not in blue light, so that it can be released throughout the cell upon illumination ^18^. However, these existing methods involve introduction of exogenous Rho-family GTPases over a background of endogenous expression. The addition of more copies of a Rho-family GTPase can lead to unnatural phenotypes, e.g. by overwhelming mechanisms that target endogenous GTPases to specific subcellular locations or by sequestering regulators such as RhoGDIs which are expressed at limited concentrations ^19^. Specifically, it has been demonstrated that expression of any Rho-family GTPase (Cdc42, Rac, and RhoA) competes with endogenous Rho-family GTPases for binding to RhoGDI-family proteins. Because RhoGDIs sequester the GDP-bound fraction of Rho-family GTPases and protect them from degradation, the introduction of exogenous Rho GTPases results in reduced levels of endogenous GTPases and constitutive activation of the remaining fraction ^20,21^ (Supplementary information, Fig. S1a-b). Thus, introducing additional GTPase molecules can produce artifactual effects on cellular behavior.

A different strategy that may produce fewer artifactual results is to use light to control upstream activators of endogenous Rho-family GTPases, rather than introducing exogenous fusion proteins of the GTPases themselves. Naturally, Dbl-family guanine nucleotide exchange factors (GEFs) constitute the primary means by which cells regulate the activity of each Rho-family GTPase ^3,22^. Various Dbl-family GEFs catalyze activation of specific Rho-family GTPases by converting the inactive GDP-bound to the active GTP-bound form, which then binds to and activate downstream effectors ^23^ (Supplementary information, Fig. S1a). Indeed, RhoA function has been optically modulated by using light-induced heterodimeric interactions such as CRY2-CIBN to increase the concentrations of a RhoGEF at the plasma membrane, where the functional subpopulation of RhoA resides ^24–26^. However, this involves overexpression of an active form of a RhoGEF throughout the cytosol, which may cause the nonspecific effect prior to light-induced membrane recruitment ^26^. Alternatively, a single-chain photoswitchable GEF can be made by fusing a GEF catalytic domain to two copies of a photodissociable variant of the green fluorescent protein Dronpa (pdDronpa) so that the active site is caged in the dark ^27,28^. Cyan light induces dissociation of the fluorescent protein domains and restoration of GEF activity, after which violet light can induce fluorescent protein domain re-association and protein re-caging. This process is reminiscent of the natural activation mechanism of some GEFs in which an upstream signal induces release of an intramolecular inhibitory interaction ^23^. By avoiding overexpression of RhoA and the resulting titration of RhoGDIs stabilizing endogenous Rho-family GTPases, this strategy may preserve more nativelike signaling states (Supplementary information, Fig. S1c).

### Development of a photoswitchable GEF for RhoA

We thus set out to construct a photocontrollable RhoA GEF. A photoswitchable Cdc42 GEF (psCdc42GEF) had been previously constructed by fusing photodissociable dimeric Dronpa (pdDronpaM) fluorescent protein domains to each end of the Dbl-homology (DH) domain of intersectin, followed by a CAAX sequence for localization to the plasma membrane ^27,28^. Baseline dimerization of pdDronpa sterically blocks the intersectin active site, while illumination causes pdDronpa dissociation and allows intersectin to bind to and activate Cdc42 ^27,28^. We investigated whether this design could be generalized by creating a photoswitchable RhoA GEF based on the RhoA-specific activator PDZRhoGEF (PRG) ^29,30^.

Previous *in vitro* findings mapped specificity determinants to only the PRG DH domain ^30^, so we first tested constructs in which photodissociable Dronpa domains (tetrameric DronpaN or dimeric pdDronpaM) were attached to both termini of the PRG DH domain, similarly to psCdc42GEF. However, HEK293A cells expressing these constructs produced a variety of responses to light, including filopodia formation and cell spreading (Supplementary information, Fig. S1a-b), suggestive of nonspecific responses from Cdc42 and Rac activation. These findings indicated that the DH domain alone in PDZRhoGEF is not sufficient to impart specificity for RhoA over other GTPases in cells, in contrast to the DH domain of the Cdc42 GE F intersectin.

We thus explored whether other elements in PRG could confer RhoA specificity in cells. While ITSN specifically bind to their substrates using only the DH domain ^31^, the crystal structure of the RhoA-PRG complex shows both the DH and PH domains of PRG contacting RhoA (Fig. 1a) ^32,33^. In addition, a “GEF switch” sequence immediately upstream of the PRG DH domain has also been suggested to interact with RhoA and contribute to its activation ^32^. Indeed, constructs in which the GEF switch sequence, DH domain, and PH domain were flanked by pdDronpaM or its higher-affinity derivative pdDronpaV (Fig. 1b) demonstrated RhoA-specific effects upon illumination (Supplementary information, Fig. S2a-b). Finally, we optimized linker lengths while testing the photodissociable pairs of pdDronpaV-pdDronpaV, pdDronpal-pdDronpal, and pdDronpal-pdDronpal N145K (Supplementary information, Fig. S2c), which form a series with decreasing affinity ^28^. pdDronpaV-PRGsDHPH-GSS_1_-pdDronpaV was most reliable in inducing cell contraction and was designated psRhoGEF (Supplementary information, Fig. S2d).

**Fig. 1.**
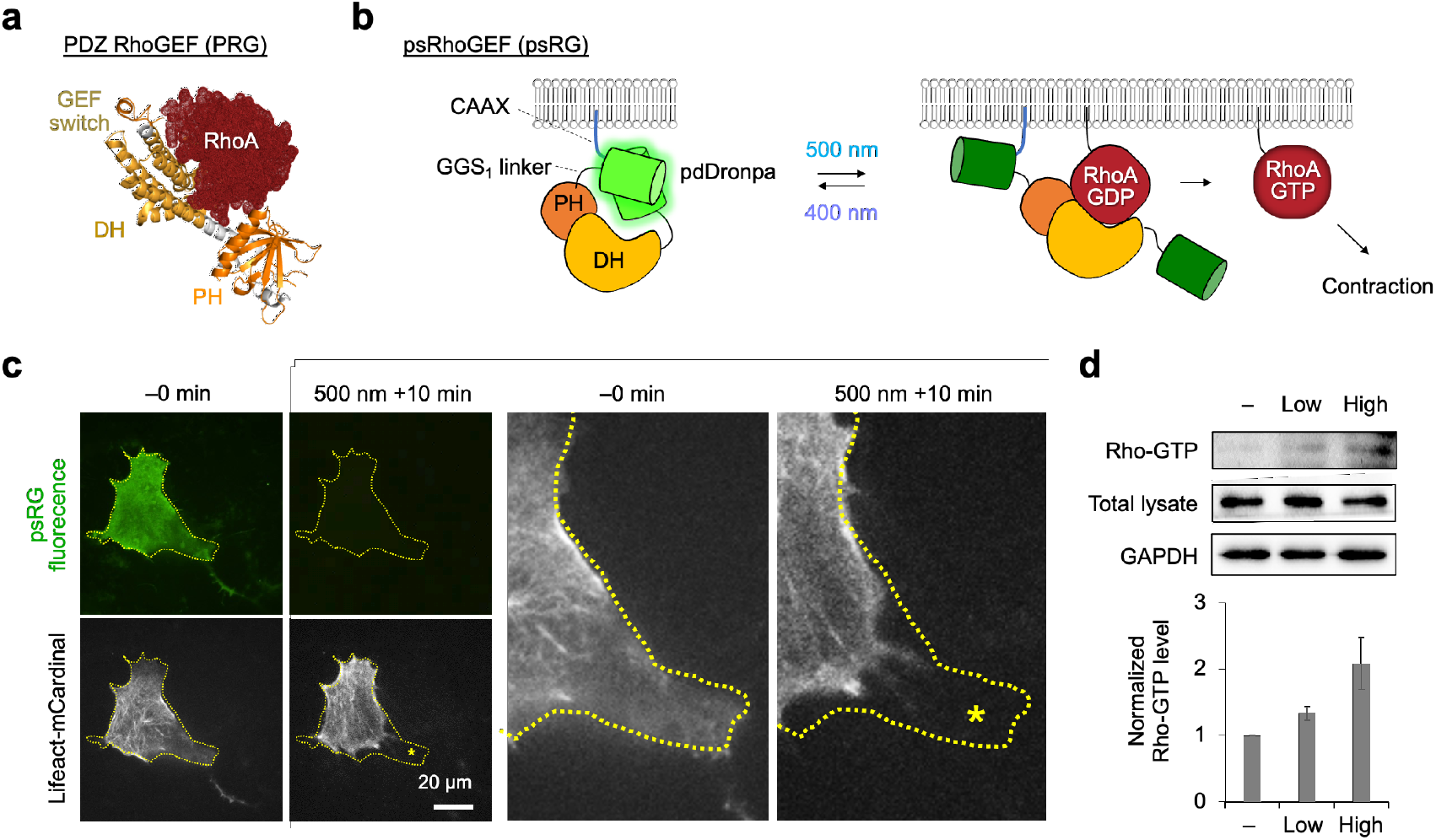
Photoswitchable RhoGEF demonstrates that acute RhoA activation induces cell retraction. (**a**) Co-crystal structure of PDZ RhoGEF (PRG) and RhoA (PDB 3T06). (**b**) Photoswitchable RhoGEF comprises the PRG RhoA-interacting regions (GEF switch, DH domain, and PH domain) fused to a pdDronpaV domain at each end with a C-terminal CAAX sequence. (**c**) Native fluorescence of pdDronpa in psRhoGEF and Lifeact-mCardinal before and after 500-nm light stimulation, showing induced cell retraction. An area was enlarged (right) to better visualize the retracted edge of the cell. A dotted line indicates original cell outline, and an asterisk indicates the region of the cell that has retracted. The experiment was repeated >10 times. See also Fig. S2. (**d**) RhoA pull-down assay confirmed different levels of active RhoA (Rho-GTP) by different light doses on psRhoGEF-expressing cells (n = 4).

psRhoGEF-induced acute RhoA activation resulted in strong cellular contraction in HEK293A cells (Fig. 1c; Supplementary information, Movie S1). We further confirmed that different light doses on psRhoGEF can induce different levels of RhoA activation (Fig. 1d). For comparison, we also tested two other optobiochemical systems for RhoA activation, OptoGEF-RhoA ^26^ and photo-recruitable (PR)-GEF ^25^. In OptoGEF-RhoA, a light-induced CRY2-CIBN interaction ^26^, while in PR-GEF, the PDZ-LOVpep interaction ^25^ (Supplementary information, Fig. S3a). Both OptoGEF-RhoA and PR-GEF mediated cell shrinkage upon illumination (Supplementary information, Fig. S3b). Thus, all three optobiochemical systems functioned effectively.

One point of difference between the systems was reversibility; after illumination was terminated, the light-induced localization of PR-GEF and OptoGEF-RhoA immediately reversed (Suppb. lementary information, Fig. S4a-b). In contrast, psRhoGEF remained in its uncaged conformation while 400-nm of illumination immediately returned it to its caged state (Supplementary information, Fig. S4c-d). In addition, psRhoGEF was less affected by expression conditions in our hands, and as its single-chain nature simplified co-expression with other constructs.

### Development of a RhoA FRET biosensor compatible with psRhoGEF

To examine RhoA activity in living cells expressing psRhoGEF, we created a RhoA FRET biosensor that could be imaged without interference by the cyan light used to activate psRhoGEF. The Large-Stokes-Shift orange fluorescent protein LSSmOrange and the far-red mKate2 are an appropriate FRET pair, which are minimally excited at the 500-nm wavelengths used to induce psRhoGEF. Thus we created a RhoA biosensor composed of LSSmOrange, the rhotekin RBD, a long linker, mKate2, and RhoA, modifying a previous design that had used cyan and yellow fluorescent proteins ^34^ (Fig. 2a-b).

**Fig. 2.**
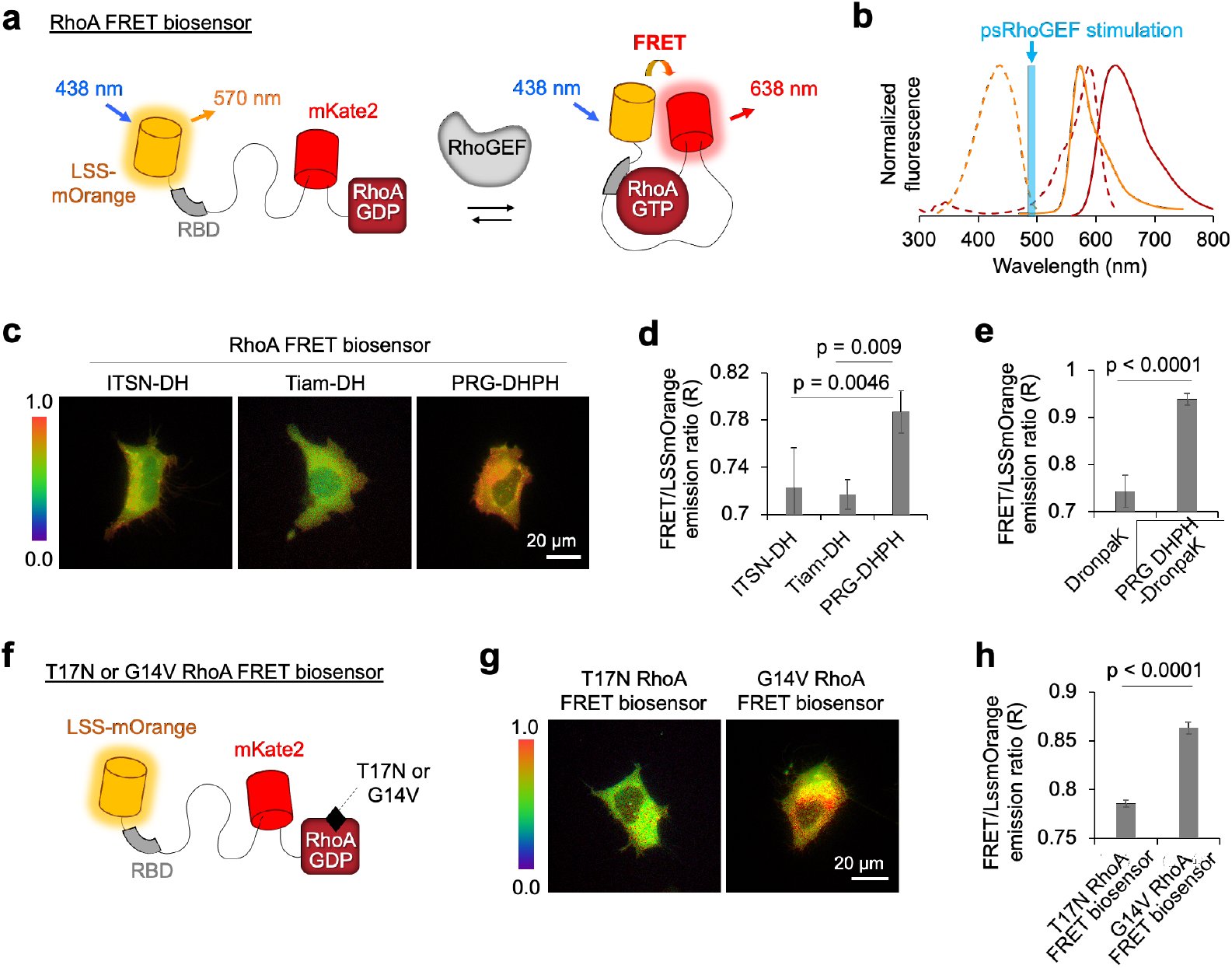
The RhoA biosensor with LSS-mOrange/mKate2 as a FRET pair. (**a**) The schematic design of a RhoA biosensor with a FRET pair, LSS-mOrange and mKate2. (**b**) Spectral profile of LSSmOrange and mKate2. Orange lines present the profile of LSSmOrange and dark red lines show the one of mKate2. The dotted and solid lines are excitation and emission spectrum, respectively. The cyan line shows peak excitation wavelength of Dronpa. (**c** and **d**) Representative images (**c**) and the average levels of FRET/LssmOrange emission ratios (**d**) in the cells expressing the RhoA biosensor together with ITSN DH, Tiam DH, or PRG DHPH (n = 40, 28, or 152, p values were calculated by Tukey’s multiple comparisons test). (**e**) The average levels of FRET/LssmOrange emission ratios in the cells expressing the RhoA biosensor together with DronpaK or DronpaK-PRG DHPH (n = 13, 18, p values were calculated by Tukey’s multiple comparisons test). (**f**) The RhoA biosensor mutants containing dominant negative (T17N) or constitutively active mutant (G14V) RhoA. (**g**) Representative images of the FRET/LSSmOrange emission ratios in the cells expressing T17N RhoA biosensor or G14V RhoA biosensor. (**h**) The average levels of the FRET/LSSmOrange emission ratios in the cells expressing dominant negative T17N RhoA or constitutively active G14V RhoA biosensor (p = 4.08e-27, by two-tailed Student’s t-test, n = 82 or 90).

To test the activity and specificity of our RhoA biosensor, we compared FRET levels in cells expressing ITSN-DH, Tiam-DH or PRG-DHPH. FRET was significantly higher in cells expressing the PRG-DHPH domains (Fig. 2c-d), indicating this sensor responds to RhoGEF with increased FRET (Fig. 2a). In contrast, the biosensor did not respond to ITSN or Tiam (Fig. 2c-d), confirming specificity for activation by RhoGEFs. We also generated constitutively active or inactive biosensors by introducing G14V or T17N mutations in the RhoA of the biosensor, respectively, and as expected, the RhoA G14V biosensor exhibited higher FRET level than the RhoA T17N biosensor (Fig. 2f-h).

We next determined whether the new RhoA biosensor can report RhoA activation by psRhoGEF. We observed that the FRET level of this biosensor in cells expressing PRG DHPH fused to monomeric DronpaK was significantly higher than in cells expressing DronpaK only (Fig. 2e), indicating that this biosensor also responds in the presence of DronpaK. We next expressed the RhoA biosensor together with psRhoGEF (Fig. 3a), and successfully observed a FRET response after illumination (Fig. 3b-c). We also found that psRhoGEF with an inactivating R868G mutation ^29,30^ failed to induce a FRET response upon illumination (Fig. 3b-c), confirming biosensor induction upon psRhoGEF photoillumination required psRhoGEF enzymatic activity. Finally, we introduced the wild-type biosensor (WT) or negative mutant T17N biosensor together with psRhoGEF. As expected, illumination caused increased FRET in the WT but not T17N biosensor (Supplementary information, Fig. S5). These results confirmed that the FRET response can report RhoA activation induced by illumination of psRhoGEF in living cells.

**Fig. 3.**
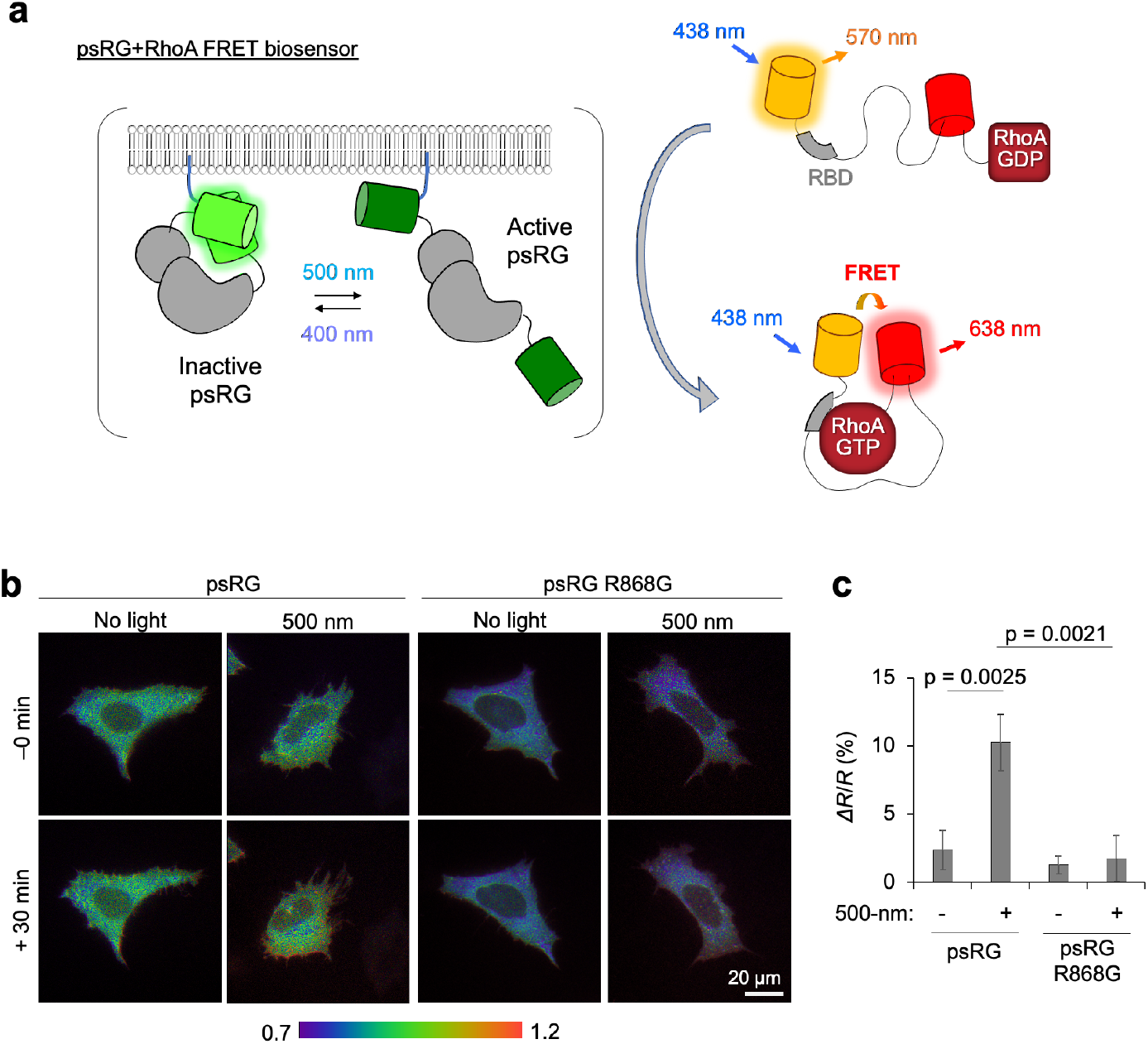
The RhoA FRET biosensor can report the RhoA activation induced by illumination on psRhoGEF. (**a**) The scheme of the FRET-based measurement of psRhoGEF-induced RhoA activation. (**b** and **c**) Representive images (**b**) and quantification (**c**) showing the changes of FRET ratios with or without light stimulation in cells co-expressing psRhoGEF or psRhoGEF R868G mutant (n = 22, 42, 14, or 21). Statistical significance is calculated by two-tailed Student’s t-test.

### Dose-dependent effects of RhoA activation on focal adhesion dynamics

RhoA has well established functions in promoting focal adhesion (FA) formation, but its potential role in FA disassembly is less clear. Following early observations that microinjection of RhoA induces FA formation in confluent fibroblasts ^1^, multiple experiments have established that RhoA and its ROCK effectors are required for FA induction by extracellular stimuli such as LPA or growth factors ^11,12,35^. If FA disassembly is considered as the reverse of FA assembly, then it might be expected that RhoA activation would inhibit FA disassembly. Indeed, chronic activation of RhoA was found to inhibit FA disassembly in fibroblasts, whereas chronic inhibition of RhoA induced FA disassembly ^36^. In addition, acute pharmacological inhibition of ROCK or myosin II, which is activated by ROCK, induces FA disassembly ^15,37^, consistent with a role for myosin in maintaining FAs. On the other hand, other experiments suggest a function for RhoA activation in FA disassembly with faster kinetics. Microinjection of RhoA induces rapid contraction of subconfluent fibroblasts ^38^. We also observed that photoinduction of psRhoGEF induced rapid shrinkage of subconfluent HEK293A cells (Fig. 1c).

We hypothesized that acute RhoA activation can induce FA disassembly during cell retraction. To test this, we expressed psRhoGEF in U87-MG astrocytoma cells, and visualized FAs with mCherry-tagged paxillin before and after illumination of 500-nm light. Indeed, we observed that acute RhoA activation induced FA disassembly at the retracted region within 30 min (Fig. 4a; Supplementary information, Movie S2). We further investigated the effect of different RhoA level on FA dynamics, taking advantage of the easy dosability of light. After applying different light intensities to psRhoGEF, different levels of RhoA activation were confirmed by the RhoA FRET biosensor (Fig. 4b).

**Fig. 4.**
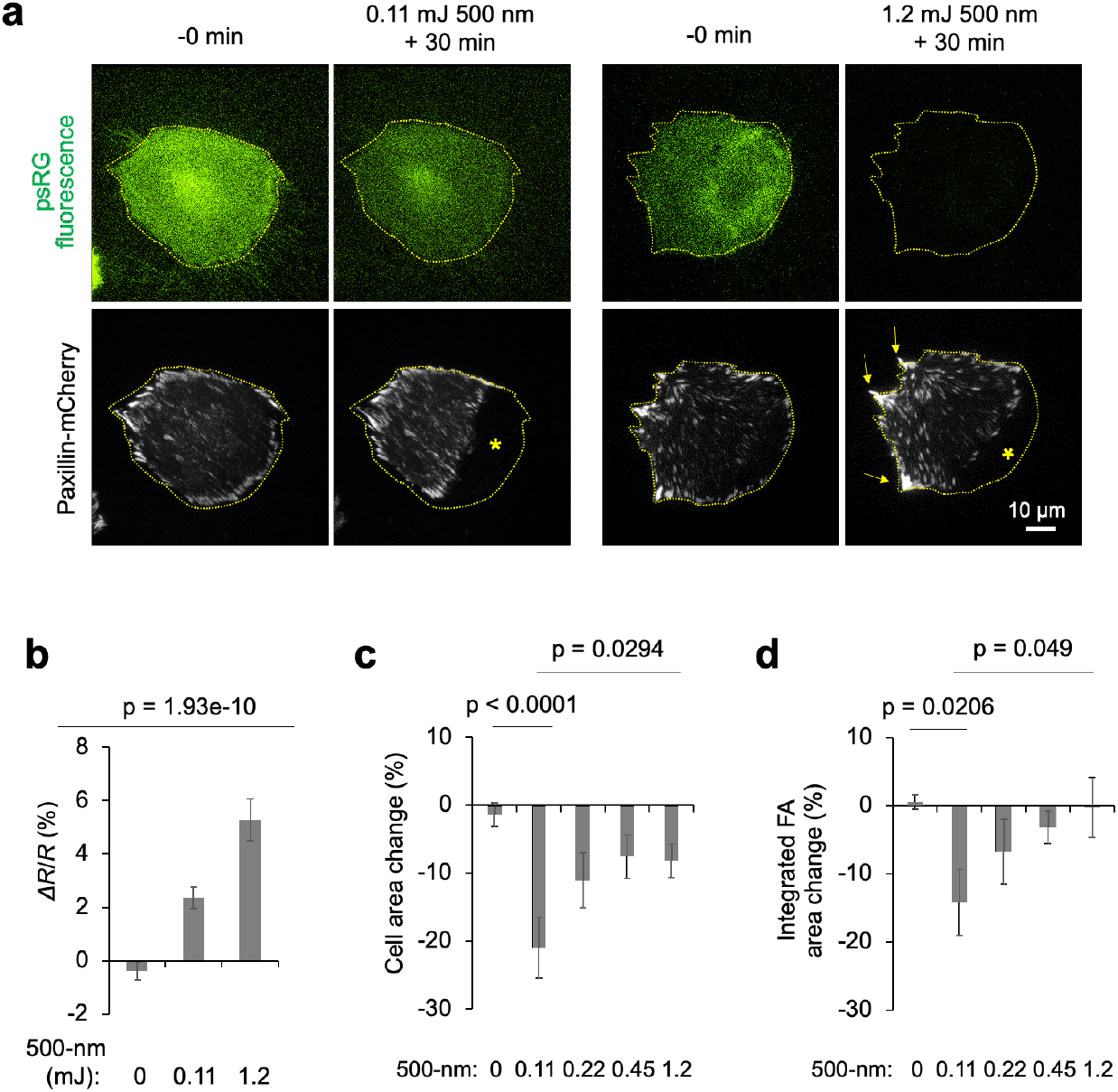
Focal adhesion responses to psRhoGEF depend on activity level. (**a**) The representative psRhoGEF and paxillin-mCherry images, before and 30 min after illumination of different light doses. (**b**) RhoA FRET ratio changes in cells expressing psRhoGEF without or with different light doses of illumination (p = 1.93e-10 by one-way ANOVA test, n = 36, 38 or 34). (**c** and **d**) The changes of cell size (**c**) and total FA area (**d**) in psRhoGEF-expressing cells in a 30-min interval without or with different doses of illumination (n = 37, 25, 18, 17 or 28). Statistical significance was calculated by Tukey’s multiple comparisons test.

Interestingly, we found unexpectedly that lower light doses favored FA disassembly and cell contraction (Fig. 4a, 4c-d). Upon 0.11 mJ of 500-nm light stimulation, we observed around 15% decrease of integrated FA area (Fig. 4d). In contrast, illumination with higher light doses (1.2 mJ) induced FA enlargement in non-retracting areas together with FA disassembly from retracting areas (Fig. 4a, right panels), so that mean integrated FA area remained unchanged (Fig. 4d). These responses are due to different RhoGEF activities (Fig. 4b) as DronpaK-expressing cells showed no changes in total cell area or total FA area at either light dose (Supplementary information, Fig. S6). Thus, we observed that psRhoGEF-induced acute RhoA activation can cause FA disassembly and cell contraction, and this cellular function of RhoA is favored under lower light doses.

### Low-amplitude RhoA signaling induces Src activation at FAs

We next investigated the downstream signaling pathways mediating this dose-dependent function of RhoA on FA dynamics. One effector of RhoA is mDia1, which mediates recruitment to FAs of Src kinase ^8^, whose activity is necessary for FA disassembly in migrating cells ^8,9^. Src phosphorylation of the FA components p130Cas and cortactin has been shown to accelerate their dissociation from FAs and be required for FA turnover in migrating cells ^39–41^.

We thus investigated the effect of different RhoA levels on Src activity at FAs after illumination of psRhoGEF-expressing cells with different light doses. Strikingly, the lower level of psRhoGEF activation induced Src activation at FAs more efficiently (Fig. 5a-b), as assessed by autophosphorylation at Tyr-416. These results imply that FA disassembly by lower RhoA activation may be due to the more efficient Src activation at FAs. To test this hypothesis, we examined the effects of perturbing Src activity on the response of FAs to low level of RhoA activation. Indeed, FA disassembly, induced by low light dose (0.11 mJ) on psRhoGEF, was completely prevented by pretreatment of a Src inhibitor PP2 (Fig. 5c).

**Fig. 5.**
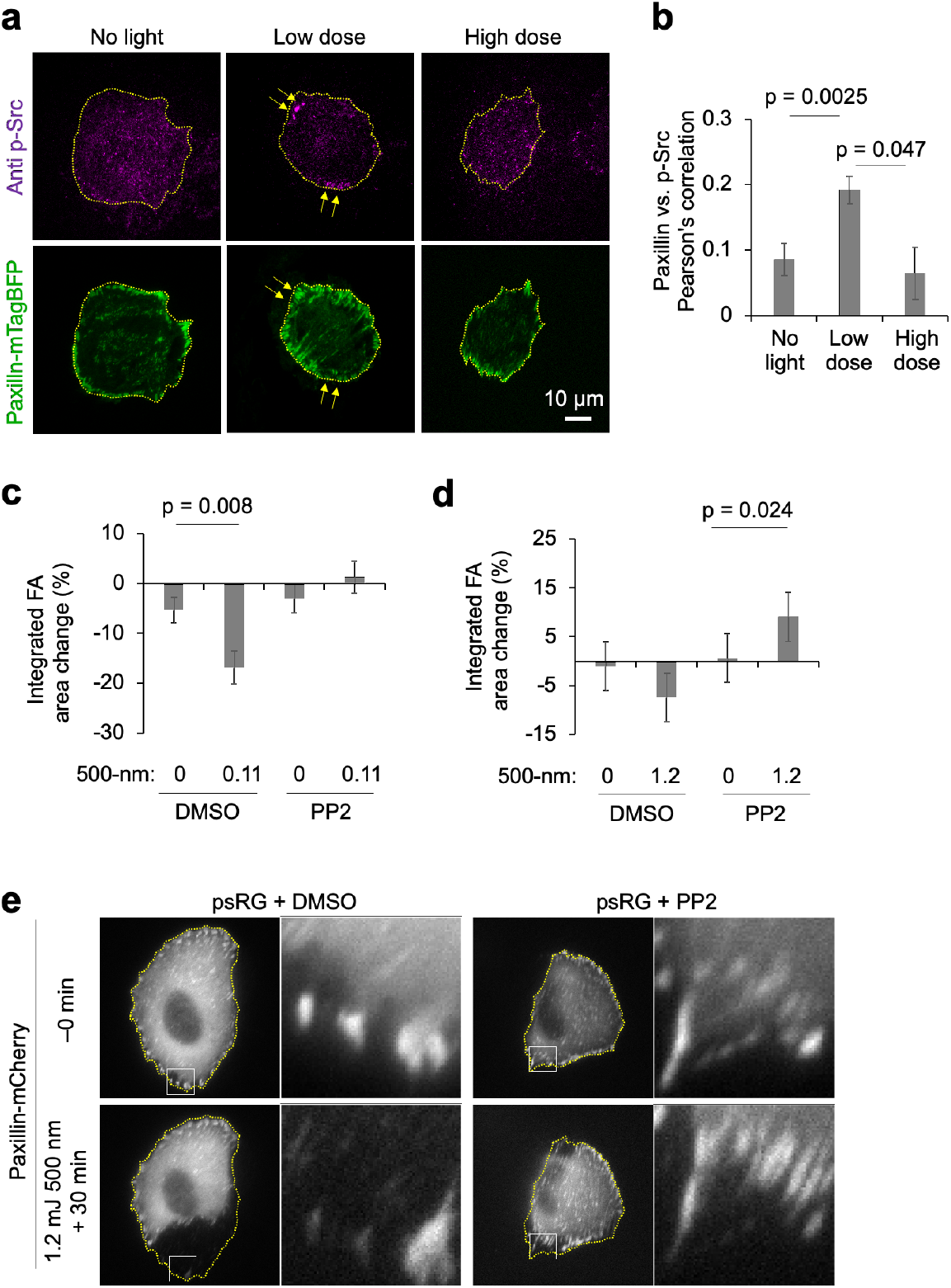
Src is activated at focal adhesions by low-level RhoA activation and mediates their disassembly. (**a**) Representative images showing differential recruitment of active p-Src at focal adhesions (labelled by paxillin-mTagBFP) upon different doses of illumination for indicated times. (**b**) Co-localization of paxillin and p-Src is estimated by the Pearson’s correlation coefficient. All p-values were calculated by two tailed Student’s t-test (n = 15, 16 or 12). (**c**) Total FA area change in psRhoGEF-expressing cells pre-treated with DMSO or 10 μM PP2 for 30 min, in a 30-min interval without or with 0.11 mJ of 500-nm light illumination (p = 0.008, two-tailed Student’s t-test, n = 71, 26, 56 or 37). (**d**) Total FA area change in psRhoGEF-expressing cells treated with DMSO or 10 μM PP2 for 30 min, in a 30-min interval without or with 1.2 mJ of 500-nm light illumination. Strong RhoA activation caused an increase in overall FA area (p = 0.024, two-tailed Student’s t-test, n = 26, 24, 25 or 23). (**e**) Representative images of FAs (labelled by paxillin-mCherry) in psRhoGEF-expressing U87-MG cells before and 30 min after 1.2 mJ of illumination, with preincubation of DMSO or 10 μM PP2 for 30 min. Peripheral FAs in the boxed areas are shown with higher magnification on the right.

We further investigated the role of Src activity on FA dynamics after high-dose (1.2 mJ) activation of psRhoGEF, with the application of the Src inhibitor PP2 or expression of dominant-negative Src K295R. Interestingly, we observed that some FAs appeared to grow inwards while still being attached to their original position at the edge of the cell and correspondingly, 1.2 mJ of illumination on psRhoGEF under PP2 incubation increased the overall FA area per cell (Fig. 5d-e; Supplementary information, Fig. S7a, Movie S3). In contrast, increasing Src activity by expression of a constitutively active Src Y527F resulted in the faster rate of cell shrinkage and focal adhesion loss in response to psRhoGEF (Supplementary information, Fig. S7b-c). These results demonstrate that Src activity is required for RhoA-induced FA disassembly, and that inhibition of Src activity can switch the response of FAs to RhoA activation from disassembly to growth.

### ROCK activity is required for both FA assembly and disassembly

Activated RhoA induce the activation of Rho-associated protein kinase ROCK. When we measured the activity of ROCK depending on light doses, and as expected, higher light doses applied on psRhoGEF induced stronger ROCK activation (Fig. 6a-b). ROCK has well established functions as a RhoA effector mediating FA growth and maturation ^16^. Indeed, FA size was significantly reduced in cells treated with the ROCK inhibitor Y-27632 (Fig. 6c-d), confirming the role of ROCK on FA.

**Fig. 6.**
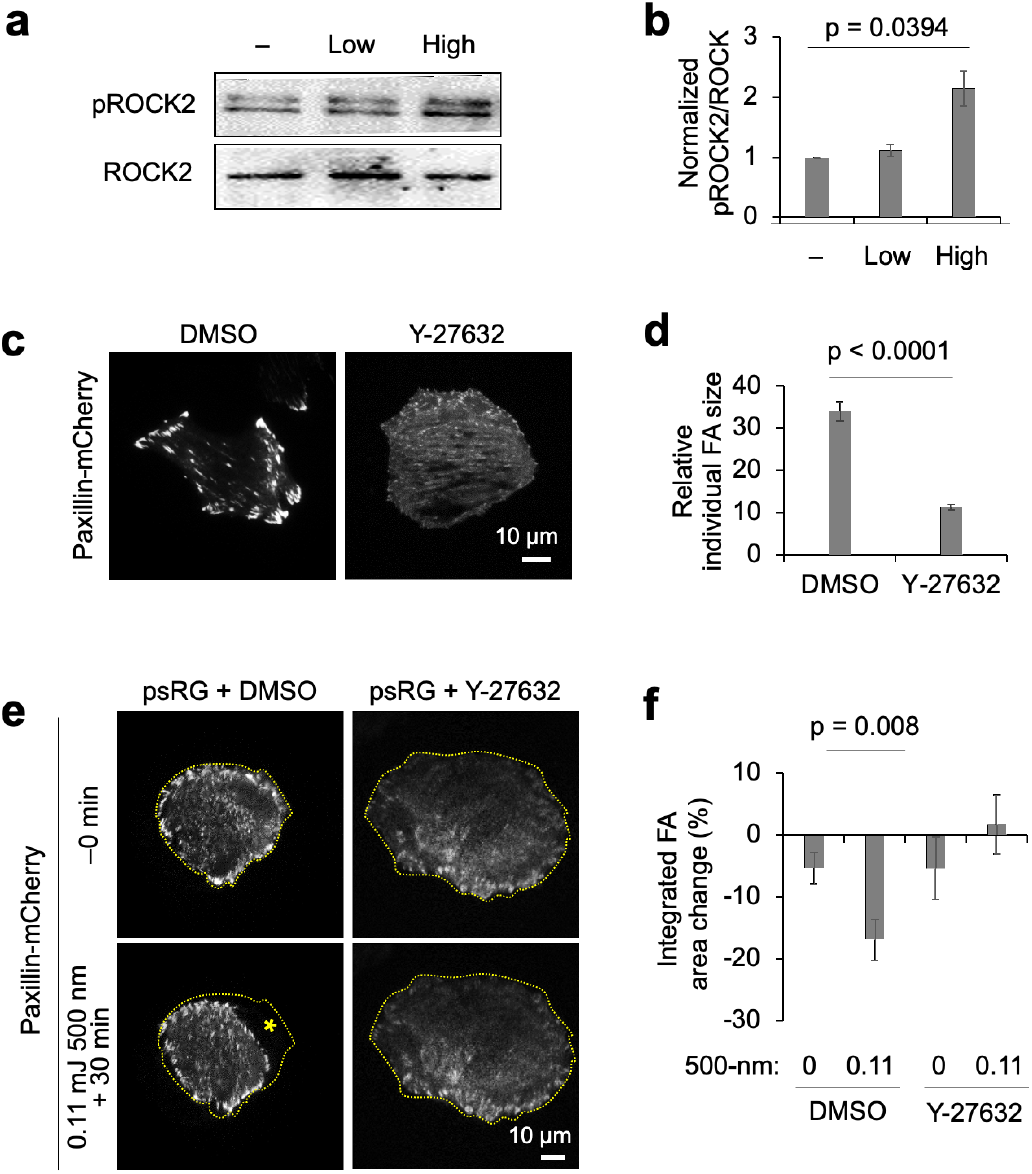
ROCK activity and actomyosin contractility are required for both FA assembly and disassembly. (**a** and **b**) (**a**) Levels of phosphorylated-ROCK2 (p-ROCK2) in U87-MG cells expressing psRhoGEF before or after different light doses. (**b**) pROCK2 was quantified relative to ROCK2. ANOVA was performed and statistical differences were calculated by post-hoc Dunnett’s test (n = 4). (**c** and **d**) Representative FA images labelled by paxillin-mCherry (**c**) and quantification of FA sizes (**d**) in psRhoGEF-expressing cells incubated with DMSO or 10 μM ROCK inhibitor Y-27632 for 30 min. Statistical differences were calculated by two tailed Student’s t-test (n=18 or 23). (**e** and **f**) Representative images (**e**) and the changes in total FA area (**f**) in psRhoGEF-expressing cells preincubated with DMSO or Y-27632 for 30 min, in a 30-min interval without or with 0.11 mJ of illumination (p = 0.008 by two tailed Student’s t-test, n = 71, 26, 23 or 21).

We next asked whether ROCK activity is required for FA disassembly that is observed at low levels of RhoA activation after 0.11 mJ of illumination on psRhoGEF. Interestingly, inhibition of ROCK by Y-27632 reduced the FA disassembly (Fig. 6e-f), indicating ROCK activity is required for RhoA-induced FA disassembly as well as FA growth.

Our results suggest that RhoA-mediated ROCK activity, in addition to Src activity at FAs, is crucial for FA disassembly. Indeed, time-lapse imaging of FA disassembly after RhoA activation revealed centripetal stretching of FAs prior to their removal (Supplementary information, Movie S3), suggesting a role for actin contractility, which is promoted by ROCK-mediated myosin II activation ^42^.

## DISCUSSION

In this study, we developed a single-chain photoswitchable activator of RhoA using a photodissociable dimeric variant of the fluorescent protein Dronpa. Using psRhoGEF to rapidly and selectively activate RhoA, we confirmed that acute RhoA activation at lower levels can induce FA disassembly, and dissected downstream pathways involved in this response. We found that both Src kinase and ROCK are required for the RhoA-induced FA disassembly, but Src phoshorylaion at FAs is more efficiently induced by lower levels of RhoA. Finally, inhibition of Src activity switches the response of FAs to RhoA activity from disassembly to enlargement. Thus RhoA activity-dependent induction of Src phosphorylation at FAs mediates the decision between the opposite outcomes of FA growth versus disassembly. In summary, our results reveal that RhoA activation can induce different FA responses in a dose-dependent manner, demonstrating that amplitude modulation of a single input can select between different outcomes of a signaling network.

Our results suggest a model for how RhoA, ROCK, and Src interact to regulate FA responses (Supplementary information, Fig. S8). Low-level RhoA activation induces strong Src phosphorylation at FAs leading to FA disassembly. In contrast, high-level RhoA activation leads to a balanced response of FA growth and disassembly, which can be biased toward FA growth by inhibition of Src. It has been suggested that RhoA has different affinity to each downstream effectors; RhoA-mDia and RhoA-ROCK complexes exhibit dissociation constants of 6 and 130 nM respectively ^43^. Thus, low concentrations of active RhoA could selectively activate mDia signaling to Src to induce FA disassembly.In contrast, high RhoA activity may preferentially activate ROCK to promote actomyosin contractility and FA growth. This model suggests that, in addition to RhoA signal amplitude, FA responses in a cell, or parts of a cell, can be modulated by the activity of other regulators of Src and ROCK.

While functions of RhoA effectors such as the ROCK and mDia in FA growth are well characterized ^16^, their roles in FA disassembly have been much less studied. For example, FAs shrink or disappear upon inhibition or depletion of ROCK or mDia ^16^. The same proteins have been implicated in FA disassembly based on more complex experiments examining rates of FA disassembly at the lagging edge of migrating cells where FA disassembly can be reliably observed ^6-8,10,16^. However, ongoing migration itself requires polarization of cells and FA assembly at the leading edge, processes that also require RhoA effectors ^6^. Thus, chronic and global manipulations of these pathways cannot unambiguously reveal direct roles for RhoA effectors in FA disassembly vs. secondary roles via affecting migration. In contrast to the use of dominant-negative or constitutively active constructs, or stimulation by extracellular stimuli, optical induction of psRhoGEF allows control of endogenous RhoA with tight temporal control and without activation of other signaling pathways. This allows immediate responses to RhoA activation to be assessed without feedback or crosstalk.

Our observation that cells can convert rheostatic RhoA signaling into opposite FA responses provides an example of how the same signal can be used to create distinct outcomes via amplitude encoding. Interestingly, RhoA may also be involved in the ability of neurons to respond to different concentrations of the chemokine SDF-1α by either enhancing or inhibiting axonal growth. While both responses require RhoA, enhancement or inhibition requires mDia or ROCK, respectively ^44^. However, in these earlier experiments, individual signaling proteins could not be specifically activated, and biological pathways could only be activated by SDF-1α application, whose downstream effects have not been comprehensively mapped. Thus, whether RhoA activity levels determine the response switch, or whether the SDF-1α receptor CXCR4 engages different signaling pathways depending on the size of the activated receptor population, could not be ascertained. In our current experiments, by directly modulating RhoA activity levels, we were able to determine that RhoA activity amplitude alone is sufficient to drive a switch between two different outcomes.

Another case of opposing responses resulting from different levels of activity of a single protein is provided by another family of small GTPases, the Ras family. Here, a low level of Ras activity induces proliferation of mammalian cells through activation of various effectors that promote protein synthesis and transcription of growth-promoting genes, whereas a high level of Ras activity caused by Ras mutation or amplification induces transcription of the cell cycle inhibitor p16 and growth arrest ^45^. However, this switch in Ras function occurs only upon mutation or amplification of Ras, a permanent genetic change, and thus cannot be dynamically regulated. In contrast, the amplitude modulation of RhoA function we observed provides a mechanism by which pathway inputs can dynamically select between alternative outputs. In other known examples of amplitude-dependent outcomes, it has not yet demonstrated that amplitude-modulation of a single intracellular signal can select between two induced outcomes (as opposed to a simple response vs. no-response decision). For example, in mammalian chondrocytes and *Xenopus* embryos, a low level of extracellular Wnt activates calcium release from internal stores, while a high level activates the β-catenin pathway ^46^. However, the mechanism behind this switch appears to be concentration-dependent utilization of different Wnt receptors, and thus response selection occurs outside rather than inside the cell. In *Drosophila* embryos, different concentrations of an epidermal growth factor homolog produce different levels of Ras activity, leading to a binary choice between transcription or no transcription ^47^, but in this case, one of the outcomes is identical to the default unstimulated state.

Finally, we proved the design of the Dronpa-based photoswitchable GEFs can be generalizable, in addition to the previously developed psCdc42GEF, by successfully developing a novel psRhoGEF. These optochemical tools should be broadly useful for investigating the functions of endogenous small Rho GTPases. For example, experiments comparing the functions of these GTPases can now be performed with a consistent set of conditions. Photoswitchable GEFs should also enable spatiotemporal control of the combination of Rho GTPases through the expression of the corresponding combination of photoswitchable GEFs. These single-chain photoswitchable GEFs enable simpler experimental designs compared to methods that utilize light-induced heterodimerization to recruit GEFs to the membrane ^24–26^. In addition, unlike optically controlled fusion proteins of Rho GTPases, these photoswitchable GEFs should avoid artifactual effects arising from the titration of a limiting number of RhoGDI molecules ^20^. We thus expect that Dronpa-based design of photoswitchable GEFs will be widely useful for investigating the functions of Rho-family GTPases with high spatiotemporal specificity and with minimal perturbation to signaling networks.

## MATERIALS AND METHODS

### DNA construction and plasmids

Plasmids were constructed by standard molecular biology methods such as polymerase chain reaction (PCR) and In-fusion cloning (Clontech). Mutations for specific amino acids were generated by overlap-extension PCR. All cloning junctions and PCR products were verified by sequencing process. The pcDNA3 vector was used for the expression in mammalian cells, and pLL3.7 vector was used for virus production. Full plasmid sequences are available upon request, and main constructs will be available in Addgene after publication.

### Cell culture and reagents

HEK293A cells were maintained in Dulbecco’s modified Eagle medium (DMEM) supplemented with 2 mM L-glutamine (GE Healthcare Life Sciences), 10% fetal bovine serum (FBS, GE Healthcare Life Sciences), 1 unit/mL penicillin (Corning), 100 g/mL streptomycin (Corning), non-essential amino acid (NEAA, Life Technologies), and 1 mM sodium pyruvate (Life Technologies). U87-MG cells were maintained in Dulbecco’s modified Eagle medium (DMEM) supplemented with 2 mM L-glutamine, 10% FBS, 1 unit/mL penicillin, 100 g/mL streptomycin, and 1 mM sodium pyruvate. Cells were cultured in a humidified 95% air, 5% CO_2_ incubator at 37°C. For the transient transfection, we used Lipofectamine 2000 (Life Technologies) or lentiviruses which were prepared by KIST Virus Facility. Src inhibitor PP2, ROCK inhibitor Y-27632 were purchased from Sigma.

### Live-cell imaging and analysis

For all imaging experiments, cover-glass-bottom dishes (SPL Life Sciences) were prepared by coating 10 μg/ml or indicated concentrations (for psRacGEF experiments) of fibronectin bovine protein (Invitrogen) for at least 2 hrs at 37°C. Cells expressing each construct were cultured on fibronectin-coated cover-glass-bottom dishes and incubated in media with 0.5% FBS overnight before imaging experiment. During imaging, cells were kept in the imaging chamber maintained with 5% CO_2_ and 37°C (Live Cell Instruments). Images were collected by a Nikon Ti-E inverted microscope with Nikon C-LHGFI HG Mercury lamp and a cooled charge-coupled device camera. Cell imaging videos were produced with MetaMorph program (Molecular Devices).

Dronpa variants were imaged with a 1/8 neutral density filter, 482/35-nm excitation filter, 506-nm dichroic mirror, 536/40-nm emission filter, and 200 ms of exposure time. mCardinal-tagged Lifeact was imaged with a 1/4 neutral density (ND 4) filter, 531/40-nm excitation filter, 562-nm dichroic mirror, 593/52-nm emission filter, and 200 ms of exposure time. Time-lapse images of Lifeact were obtained every one-minute, for 10 min before photoswitch and for 50 min after photoswitch. Dronpa variants were photoswitched off by illumination on a 100× oil objective (numberical aperture 1.45, working distance 0.13 mm, Nikon) using the same excitation and dichroic filters as for imaging but without a neutral density filter for 30 seconds. For different light doses, Dronpa was photoswitched by illumination using a 100x oil objective without a neutral density filter (full dose) or 1/2, 1/4, or 1/8 neutral density (50%, 25%, or 12.5%) for 30 seconds.

Focal adhesion (FA) area for each cell at each time point was automatically calculated using the Focal Adhesion Analysis Server (FAAS) ^48^. Cells were categorized as showing FA growth or FA shrinkage if total FA area change during the experiment was higher than the standard deviation in the non-stimulated population.

For the FRET imaging, mKate emission from FRET was imaged by 438/24-nm excitation filter, 593-nm dichroic mirror, 641/75-nm emission filter for 200 ms of exposure time. LSSmOrange emission image was collected with 438/24-nm excitation filter, 562-nm dichroic mirror, 593/40-nm emission filter, for 200 ms of exposure time. After background-subtraction, the pixel-by-pixel ratio images of FRET/LSSmOrange were calculated by the NIS program. The average FRET/LSSmOrange ratio in total cell area (R) or its relative change after stimulation (ΔR/R) were calculated for the statistical analysis of FRET responses.

Total internal reflection fluorescence (TIRF) images of mCherry-tagged paxillin before and after psRhoGEF activation were acquired under Nikon Ti-E inverted microscope equipped with fiber-coupled 488-nm and 561-nm lasers to excite Dronpa and mCherry, respectively. NIS-elements software was used for image acquisition and analysis.

### Rho GTPase pull-down assay

U87-MG cells stably infected by lentviral psRhoGEF were lysed with 1X MLB buffer (EMD Millipore) containing 25 mM, 1mM sodium orthovanadate, and protease inhibitor cocktail tablet (Roche). All samples were quantified and agitated overnight at 4°C with 23 μl of Rhotekin RBD agarose bead (EMD Millipore). Equal amounts of protein were subject to SDS-PAGE and blotted with anti-Rho antibody (3 μg/mL, #05-778, EMD Millipore) or anti-GAPDH antibody (1:1000 dilution, #SC47724, Santa Cruz Biotechnology). We developed western blot membranes with enhanced chemiluminescence (ECL) solution and images were captured with a luminescent image analyzer ChemiDoc (Bio-Rad, USA) or Amersham Imagequant 800 (Cytiva, USA).

### Western blotting

U87-MG cells infected by psRhoGEF containing lentiviruses were cultured for around 30 hrs and then starved in 0.5% FBS media for overnight. These cells were lysed with cell lysis buffer (Cell signaling) containing 1 mM PMSF, 5 mM NaF, and protease inhibitor cocktail (Cytoskeleton). Equal amounts of protein were subject to SDS-PAGE and blotted with polyclonal anti-phosphorylated ROCK2 (Ser1366) antibody (1.3 μg/mL, #ab228008, Abcam) or polyclonal anti-ROCK2 antibody (1:1000 dilution, #8236, Cell Signaling Technology). We developed western blot membranes with enhanced chemiluminescence (ECL) solution and images were captured with a luminescent image analyzer ChemiDoc (Bio-Rad, USA).

### Immunostaining

Paxillin-mTagBFP and psRhoGEF co-expressed cells were fixed with 4% paraformaldehyde for 10 min, and permeabilized with 0.1% triton X-100 for 15 min. Cells were incubated in 5% BSA in PBS for 1 hr, and then with rabbit anti-phospho Src Tyr416 (0.69 μg/mL, #2101, Cell Signaling) for overnight at 4°C. After three times of washing with PBS, cells were incubated with goat anti-rabbit antibody conjugated to Alexa-fluor 594 (20 μg/mL, #A-11037, Thermo Fisher Scientific) for 1 hr. After three times of washing with PBS for 10 min each, the stained cells were mounted and observed under a TIRF microscopy. For quantifying of p-Src at focal adhesions, the TIRF images of p-Src and paxillin-mTagBFP of the same cell were applied to the Pearson’s correlation test via NIS program (Nikon).

### Statistical tests

P values were calculated using two-tailed Student’s t-test, the Dunnett’s multiple comparisons test, one-way ANOVA test or Tukey’s multiple comparisons test (GraphPad Prism 8) for continuous variables, following confirmation of normality calculated by the Shapiro-Wilk test calculator (Statistics Kingdom), or using Fisher’s exact test calculator (Social Science Statistics, 2018) for categorical variables.

## ACKNOWLEDGEMENTS

This work is supported by KIST Institutional Grant 2E30963, Brain Research Program through the National Research Foundation of Korea (2017M3C7A1043842), Samsung Research Funding & Incubation Center of Samsung Electronics under Project Number SRFC-TC2003-02 (J.S.), the Stanford Department of Bioengineering (A.L.-R.) and a NIH Pioneer Award (L.N., X.X.Z., M.Z.L.).

## AUTHOR CONTRIBUTIONS

J.S. and M.Z.L. designed research; J.S., J.J., H.N.L, L.N., H.R., X.X.Z., H.C., Y.W.L, A.L.-R. performed experiments; J.S., M.Z.L., J.J, L.N., C. J. analyzed data; J.S. and M.Z.L. wrote the manuscript.

## ADDITIONAL INFORMATION

**Supplementary information** is aviable.

**Competing interests: The authors declare no conflicts of interest.**

## REFERENCES

1 Nobes, C. D. & Hall, A. Rho, rac, and cdc42 GTPases regulate the assembly of multimolecular focal complexes associated with actin stress fibers, lamellipodia, and filopodia. Cell 81, 53–62 (1995).

2 Heasman, S. J. & Ridley, A. J. Multiple roles for RhoA during T cell transendothelial migration. Small GTPases 1, 174–179, doi:10.4161/sgtp.1.3.14724 (2010).

3 Machacek, M. et al. Coordination of Rho GTPase activities during cell protrusion. Nature 461, 99–103, doi:10.1038/nature08242 (2009).

4 Pertz, O., Hodgson, L., Klemke, R. L. & Hahn, K. M. Spatiotemporal dynamics of RhoA activity in migrating cells. Nature 440, 1069–1072, doi:10.1038/nature04665 (2006).

5 Wong, K., Pertz, O., Hahn, K. & Bourne, H. Neutrophil polarization: spatiotemporal dynamics of RhoA activity support a self-organizing mechanism. Proc Natl Acad Sci U S A 103, 3639–3644, doi:10.1073/pnas.0600092103 (2006).

6 Parsons, J. T., Horwitz, A. R. & Schwartz, M. A. Cell adhesion: integrating cytoskeletal dynamics and cellular tension. Nat Rev Mol Cell Biol 11, 633–643, doi:10.1038/nrm2957 (2010).

7 Lock, F. E., Ryan, K. R., Poulter, N. S., Parsons, M. & Hotchin, N. A. Differential regulation of adhesion complex turnover by ROCK1 and ROCK2. PLoS One 7, e31423, doi:10.1371/journal.pone.0031423 (2012).

8 Yamana, N. et al. The Rho-mDia1 pathway regulates cell polarity and focal adhesion turnover in migrating cells through mobilizing Apc and c-Src. Mol Cell Biol 26, 6844–6858, doi:10.1128/MCB.00283-06 (2006).

9 Carragher, N. O. & Frame, M. C. Focal adhesion and actin dynamics: a place where kinases and proteases meet to promote invasion. Trends Cell Biol 14, 241–249, doi:10.1016/j.tcb.2004.03.011 (2004).

10 Webb, D. J. et al. FAK-Src signalling through paxillin, ERK and MLCK regulates adhesion disassembly. Nat Cell Biol 6, 154–161, doi:10.1038/ncb1094 (2004).

11 Amano, M. et al. Formation of actin stress fibers and focal adhesions enhanced by Rho-kinase. Science 275, 1308–1311 (1997).

12 Oakes, P. W., Beckham, Y., Stricker, J. & Gardel, M. L. Tension is required but not sufficient for focal adhesion maturation without a stress fiber template. J Cell Biol 196, 363–374, doi:10.1083/jcb.201107042 (2012).

13 Meenderink, L. M. et al. P130Cas Src-binding and substrate domains have distinct roles in sustaining focal adhesion disassembly and promoting cell migration. PLoS One 5, e13412, doi:10.1371/journal.pone.0013412 (2010).

14 Stricker, J., Beckham, Y., Davidson, M. W. & Gardel, M. L. Myosin II-mediated focal adhesion maturation is tension insensitive. PLoS One 8, e70652, doi:10.1371/journal.pone.0070652 (2013).

15 Wolfenson, H., Bershadsky, A., Henis, Y. I. & Geiger, B. Actomyosin-generated tension controls the molecular kinetics of focal adhesions. J Cell Sci 124, 1425–1432, doi:10.1242/jcs.077388 (2011).

16 Burridge, K. & Guilluy, C. Focal adhesions, stress fibers and mechanical tension. Exp Cell Res 343, 14–20, doi:10.1016/j.yexcr.2015.10.029 (2016).

17 Bugaj, L. J., Choksi, A. T., Mesuda, C. K., Kane, R. S. & Schaffer, D. V. Optogenetic protein clustering and signaling activation in mammalian cells. Nat Methods 10, 249–252, doi:10.1038/nmeth.2360 (2013).

18 Wang, H. et al. LOVTRAP: an optogenetic system for photoinduced protein dissociation. Nat Methods 13, 755–758, doi:10.1038/nmeth.3926 (2016).

19 Michaelson, D. et al. Differential localization of Rho GTPases in live cells: regulation by hypervariable regions and RhoGDI binding. J Cell Biol 152, 111–126 (2001).

20 Boulter, E. & Garcia-Mata, R. RhoGDI: A rheostat for the Rho switch. Small GTPases 1, 65–68, doi:10.4161/sgtp.1.1.12990 (2010).

21 Bozza, W. P., Zhang, Y., Hallett, K., Rivera Rosado, L. A. & Zhang, B. RhoGDI deficiency induces constitutive activation of Rho GTPases and COX-2 pathways in association with breast cancer progression. Oncotarget 6, 32723–32736, doi:10.18632/oncotarget.5416 (2015).

22 Luo, L. Rho GTPases in neuronal morphogenesis. Nat Rev Neurosci 1, 173–180, doi:10.1038/35044547 (2000).

23 Rossman, K. L., Der, C. J. & Sondek, J. GEF means go: turning on RHO GTPases with guanine nucleotide-exchange factors. Nat Rev Mol Cell Biol 6, 167–180, doi:10.1038/nrm1587 (2005).

24 O’Neill, P. R., Kalyanaraman, V. & Gautam, N. Subcellular optogenetic activation of Cdc42 controls local and distal signaling to drive immune cell migration. Mol Biol Cell 27, 1442–1450, doi:10.1091/mbc.E15-12-0832 (2016).

25 Wagner, E. & Glotzer, M. Local RhoA activation induces cytokinetic furrows independent of spindle position and cell cycle stage. J Cell Biol 213, 641–649, doi:10.1083/jcb.201603025 (2016).

26 Valon, L., Marin-Llaurado, A., Wyatt, T., Charras, G. & Trepat, X. Optogenetic control of cellular forces and mechanotransduction. Nat Commun 8, 14396, doi:10.1038/ncomms14396 (2017).

27 Zhou, X. X., Chung, H. K., Lam, A. J. & Lin, M. Z. Optical control of protein activity by fluorescent protein domains. Science 338, 810–814, doi:10.1126/science.1226854 (2012).

28 Zhou, X. X., Fan, L. Z., Li, P., Shen, K. & Lin, M. Z. Optical control of cell signaling by single-chain photoswitchable kinases. Science 355, 836–842, doi:10.1126/science.aah3605 (2017).

29 Jaiswal, M. et al. Mechanistic insights into specificity, activity, and regulatory elements of the regulator of G-protein signaling (RGS)-containing Rho-specific guanine nucleotide exchange factors (GEFs) p115, PDZ-RhoGEF (PRG), and leukemia-associated RhoGEF (LARG). J Biol Chem 286, 18202–18212, doi:10.1074/jbc.M111.226431 (2011).

30 Oleksy, A., Opalinski, L., Derewenda, U., Derewenda, Z. S. & Otlewski, J. The molecular basis of RhoA specificity in the guanine nucleotide exchange factor PDZ-RhoGEF. J Biol Chem 281, 32891–32897, doi:10.1074/jbc.M606220200 (2006).

31 Snyder, J. T. et al. Structural basis for the selective activation of Rho GTPases by Dbl exchange factors. Nat Struct Biol 9, 468–475, doi:10.1038/nsb796 (2002).

32 Bielnicki, J. A. et al. Insights into the molecular activation mechanism of the RhoA-specific guanine nucleotide exchange factor, PDZRhoGEF. J Biol Chem 286, 35163–35175, doi:10.1074/jbc.M111.270918 (2011).

33 Kristelly, R., Gao, G. & Tesmer, J. J. Structural determinants of RhoA binding and nucleotide exchange in leukemia-associated Rho guanine-nucleotide exchange factor. J Biol Chem 279, 47352–47362, doi:10.1074/jbc.M406056200 (2004).

34 Fritz, R. D. et al. A versatile toolkit to produce sensitive FRET biosensors to visualize signaling in time and space. Sci Signal 6, rs12, doi:10.1126/scisignal.2004135 (2013).

35 Ridley, A. J., Paterson, H. F., Johnston, C. L., Diekmann, D. & Hall, A. The small GTP-binding protein rac regulates growth factor-induced membrane ruffling. Cell 70, 401–410 (1992).

36 Ren, X. D. et al. Focal adhesion kinase suppresses Rho activity to promote focal adhesion turnover. J Cell Sci 113 (Pt 20), 3673–3678 (2000).

37 Seong, J. et al. Distinct biophysical mechanisms of focal adhesion kinase mechanoactivation by different extracellular matrix proteins. Proc Natl Acad Sci U S A 110, 19372–19377, doi:10.1073/pnas.1307405110 (2013).

38 Paterson, H. F. et al. Microinjection of recombinant p21rho induces rapid changes in cell morphology. J Cell Biol 111, 1001–1007 (1990).

39 Machiyama, H. et al. Displacement of p130Cas from focal adhesions links actomyosin contraction to cell migration. J Cell Sci 127, 3440–3450, doi:10.1242/jcs.143438 (2014).

40 Sawada, Y. et al. Force sensing by mechanical extension of the Src family kinase substrate p130Cas. Cell 127, 1015–1026, doi:10.1016/j.cell.2006.09.044 (2006).

41 Wang, W., Liu, Y. & Liao, K. Tyrosine phosphorylation of cortactin by the FAK-Src complex at focal adhesions regulates cell motility. BMC Cell Biol 12, 49, doi:10.1186/1471-2121-12-49 (2011).

42 Amano, M., Nakayama, M. & Kaibuchi, K. Rho-kinase/ROCK: A key regulator of the cytoskeleton and cell polarity. Cytoskeleton (Hoboken) 67, 545–554, doi:10.1002/cm.20472 (2010).

43 Narumiya, S., Tanji, M. & Ishizaki, T. Rho signaling, ROCK and mDia1, in transformation, metastasis and invasion. Cancer Metastasis Rev 28, 65–76, doi:10.1007/s10555-008-9170-7 (2009).

44 Arakawa, Y. et al. Control of axon elongation via an SDF-1alpha/Rho/mDia pathway in cultured cerebellar granule neurons. J Cell Biol 161, 381–391, doi:10.1083/jcb.200210149 (2003).

45 Dimauro, T. & David, G. Ras-induced senescence and its physiological relevance in cancer. Curr Cancer Drug Targets 10, 869–876 (2010).

46 Kestler, H. A. & Kuhl, M. Generating a Wnt switch: it’s all about the right dosage. J Cell Biol 193, 431–433, doi:10.1083/jcb.201103167 (2011).

47 Sagner, A. & Briscoe, J. Morphogen interpretation: concentration, time, competence, and signaling dynamics. Wiley Interdiscip Rev Dev Biol 6, doi:10.1002/wdev.271 (2017).

48 Berginski, M. E. & Gomez, S. M. The Focal Adhesion Analysis Server: a web tool for analyzing focal adhesion dynamics. F1000Res 2, 68, doi:10.12688/f1000research.2-68.v1 (2013).

